# Complementary modes of resistance to *EGFR* TKI in lung adenocarcinoma through MAPK activation and cellular plasticity

**DOI:** 10.1101/2025.05.07.652714

**Authors:** Matthew Zatzman, Alvaro Quintanal-Villalonga, Sohrab Salehi, Nicholas Ceglia, Jake June-Koo Lee, Amanda N. Pupo, Christina J. Falcon, Nicole Rusk, Ignas Masilionis, Ojasvi Chaudhary, Parvathy Manoj, Ronan Chaligne, Andrew McPherson, Charles M. Rudin, Sohrab P. Shah, Helena A. Yu

## Abstract

*EGFR*-mutant lung adenocarcinoma (LUAD) represents 20% of all non-small cell lung carcinomas, with most patients presenting with incurable metastatic disease. Treatment with mutant-selective *EGFR* tyrosine kinase inhibitors (TKIs) results in initial tumor reduction, yet nearly all patients eventually relapse. The mechanisms driving drug resistance are incompletely understood, creating significant barriers to curing metastatic disease. We integrated clinical genomic and single-nuclei RNA (snRNA) sequencing from a cohort of 62 *EGFR*-mutant LUAD patients treated with the third generation *EGFR* TKI, osimertinib, and compared treatment-naïve (TN), minimal residual disease (MRD), and progressive disease (PD) tumors. We found that disease progression is associated with a marked decrease in alveolar lineage fidelity, coincident with reduced MAPK signaling and adenocarcinoma identity. PD tumors with sustained MAPK pathway activity, such as those with *EGFR* or *MET* amplifications, tended to retain adenocarcinoma identity. In contrast, MAPK-low tumors were more likely to undergo histological transformation to squamous or neuroendocrine lineages. Remarkably, we observed rare tumor cell populations prior to treatment that were poorly differentiated, in some cases with neuroendocrine or squamous features. At progression, these histologically divergent tumor cells increased in prevalence, both in cases with overt histological transformation, and in others with sub-clinical histological plasticity. These findings suggest that pre-existing capacity for histologic plasticity may be a substrate for therapy induced selection. Taken together, our results illuminate genomically encoded MAPK signaling and lineage plasticity as complementary mechanisms of acquired resistance to *EGFR* TKI in lung adenocarcinoma.

## INTRODUCTION

Lung adenocarcinoma (LUAD) is characterized by an epithelial alveolar phenotype closely related to alveolar type 2 (AT2) cells – thought to be the cell type of origin^1,2^. Epidermal Growth Factor Receptor *(EGFR)*-mutant LUAD represents approximately 20% of all non-small cell lung carcinomas (NSCLC), and up to 50% in east Asian countries^3^. These tumors are driven by mutation-induced constitutive activation of *EGFR*, leading to dysregulated induction of downstream pathways such as mitogen-activated protein kinase (MAPK) signaling^4^. While treatment with *EGFR* tyrosine kinase inhibitors (TKIs), such as osimertinib, initially results in marked tumor shrinkage, it typically fails to completely eradicate disease, resulting in drug resistance in nearly all cases. Upon disease progression, treatment options are limited to chemotherapies, which have minimal efficacy and increased toxicity, underscoring the need for improved therapeutic interventions.

Resistance to *EGFR* TKIs is characterized by both genomic and non-genomic adaptation. Secondary genomic alterations, detected in up to 50% of relapsed patients, typically converge on MAPK signaling pathways^5–7^. These alterations are classified as on-target or off-target depending on whether they occur in *EGFR* (e.g. C797X or amplification) or other oncogenes respectively (e.g. *MET*, *ERBB2*, *KRAS* and others) but in general result in sustained MAPK signaling. Non-genetic adaptation is also implicated in the emergence of drug resistance. In the majority of cases, a clinically detectable tumor mass remains during treatment, known as minimal residual disease (MRD). These persister tumor cells are characterized by a stem-like, slow proliferating phenotype which may confer adaptive capacity, eventually overcoming treatment-induced fitness barriers to further expansion^8–12^. Epithelial-to-mesenchymal transition (EMT), another phenomenon initially described as a key step during the metastatic process and thought to be non-genomically driven, is also associated with TKI resistance^13^. More rarely, tumors can circumvent *EGFR* inhibition by undergoing histological transformation to either lung squamous carcinoma (LUSC) or small cell lung carcinoma (SCLC), occurring in approximately 9 and 14% of osimertinib-relapsed tumors, respectively^14–16^.

Tumor tissue is rarely collected during treatment, or at the time of progression, with most available datasets including only genomic characterization derived from bulk tissue sequencing^5^, or a limited sampling of cells from EGFR mutant cases^17^. As a result, our understanding of: i) the tumor cell intrinsic transcriptional adaptations that accompany drug resistance in patients, ii) whether they exist prior to treatment, and iii) the complex interplay between genomic and non-genomic mechanisms that might govern these shifts, remains incomplete. Furthermore, it is unclear to what extent the MRD phenotype exists prior to therapy exposure, and if it persists and/or expands into progressive disease. Here, we leveraged combined single-cell transcriptomic and clinical genomic profiling in a cohort of *EGFR*-mutant LUAD patients sampled at distinct disease stages, uncovering distinct modes of drug resistance defined by alveolar cell identity and sustained MAPK signaling.

## RESULTS

### Multimodal profiling of osimertinib treated *EGFR* mutant lung cancers

Tumor tissue was collected from 62 unmatched patients with *EGFR*-mutant LUAD, including osimertinib treatment-naive (TN; n=26), minimal residual disease (MRD) samples taken on osimertinib (n=6), and progressive disease (PD) samples (n=30; **Fig. 1**). Three TN patients had been exposed to treatments other than osimertinib prior to sample collection, including erlotinib in TN1, and chemo-IO for TN14 and TN16, who had a concurrent *KRAS* G12C stage IV LUAD, and recurrent triple negative breast cancer, respectively. Tumor stage and tissue sites varied across the treatment timepoints, with TN samples consisting mainly of early stage (I-II) disease sampled from primary tumor lung sites, whereas MRD and PD samples were exclusively stage IV tumor specimens taken from either primary lung (n=13), or metastatic sites including intrapulmonary metastases (locoregional, n=3), lymph-nodes (n=2), bone and soft tissue (n=7), liver (n=6), brain (n=3) and pleura (n=2). Samples were profiled using single -nuclei RNA sequencing (snRNAseq; n=62), clinical tumor-normal DNA sequencing using a 468 cancer-gene panel (MSK-IMPACT^18^, n=57) and by histopathological evaluation **(Fig. S1, Table S1)**. *EGFR* mutation status was confirmed in all patients by clinical testing and/or MSK-IMPACT sequencing, with exon 19 alterations being the most common, followed by L858R. Four patients had T790M alterations, which in three cases was likely related to prior erlotinib exposure (TN1, PD2, PD3). *EGFR* on-target alterations were identified in 11/30 PD patients, with genomic amplifications being the most common (n=10), and one patient (PD26) harboring a *EGFR* C797S mutation. As expected given their later stage, MRD and PD samples had a significant increase in both point mutation and copy number burden compared to earlier stage TN tumors **(Fig. S2A-E)**. Consistent with this, PD tumors were enriched for mutations in *TP53*, *RB1* and whole-genome doubling events **(Fig. S2F-G)**.

**Figure 1.**
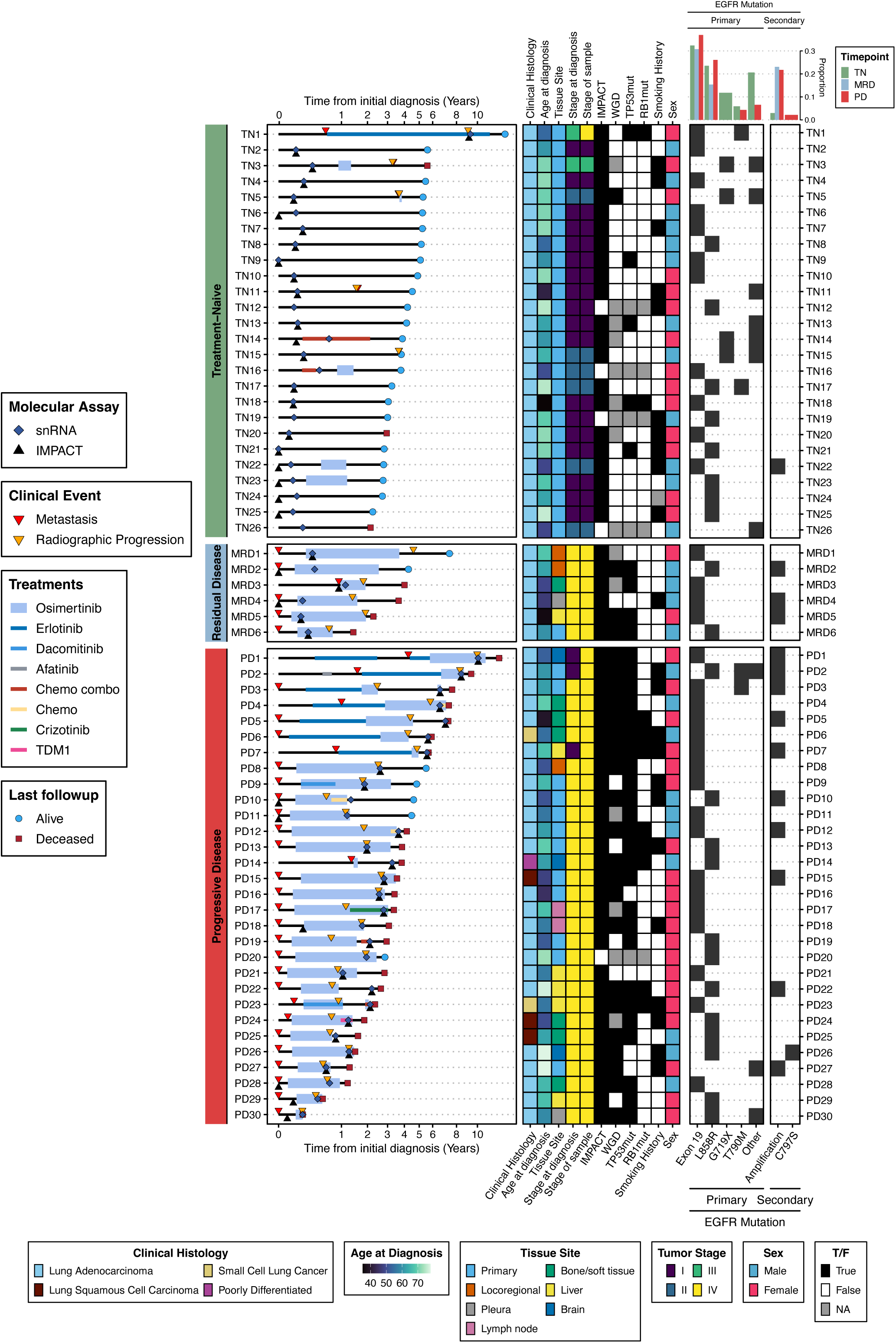
Multimodal profiling of osimertinib treated *EGFR* mutant lung cancers. Patient swimmer’s lane plot depicting treatment course, clinical events, and assay collection times for each patient (left), clinical metadata (middle), *EGFR* mutation types (right), and *EGFR* mutation type proportions (top-right).

Using snRNAseq, we measured gene expression profiles in 385,770 cells across our cohort, defining six broad cell type compartments: (1) tumor and epithelial cells; (2) T and NK cells; (3) myeloid cells; (4) fibroblasts; (5) endothelial cells and (6) B and plasma cells (**Fig. 2A-D, Fig. S3A-B**). Malignant and non-malignant epithelial cells were distinguished on the basis of inferred copy number profiles from snRNA (**Fig. S4A, Methods**). Tumor cellularity varied across samples (range 0-95%), with 6 samples harboring no identifiable tumor cell population (**Fig. S4B)**. While TN and PD samples showed no difference in terms of tumor cellularity, MRD samples exhibited significantly lower tumor cell content, consistent with the effect of ongoing treatment (**Fig. 2E, Fig. S4C**). Tumor cellularity was relatively consistent across tissue sites and procedure types, though pleural samples and lobectomies tended to yield fewer tumor cells overall (**Fig. S4D-E**).

**Figure 2.**
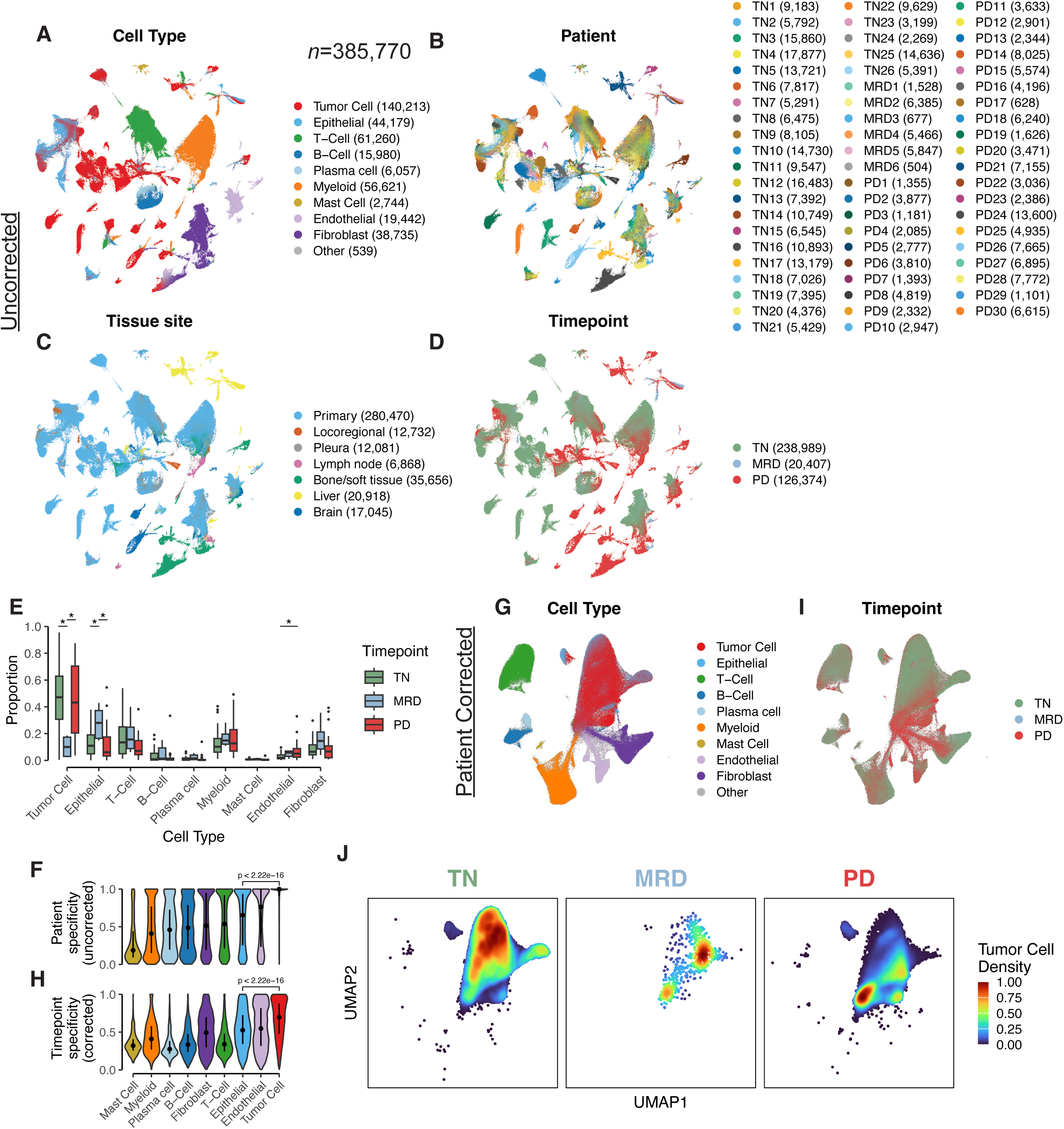
Tumor cell intrinsic phenotypic shifts accompany disease progression. **A-D.** UMAP plot of cells prior to batch correction, colored by cell type **(A)**, patient (B), tissue site **(C)** and treatment timepoint **(D)**. **E.** Boxplot of major cell type proportions across treatment timepoints. Wilcoxon rank-sum test significance is annotated as ‘*’: p<0.05. **F.** Patient specificity scores for each cell type on batch uncorrected data. Dot and linerange indicate median and IQR. **G,I.** UMAP plot of cells after batch correction, colored by cell type and timepoint. **H.** Timepoint specificity scores for each cell type on batch corrected data. Dot and linerange indicate median and IQR. **J.** Density of tumor cells from each treatment timepoint overlaid on the batch corrected UMAP.

Tumor cells exhibited a significantly higher degree of patient- **(Fig. 2B,F),** site- (**Fig. 2C)** and treatment timepoint specificity (**Fig. 2D)** compared to non-malignant epithelium. We regressed out patient specific effects to better understand phenotypic shifts as a function of disease state (**Fig. 2G, Fig. S5A-B, Methods**). Post-correction, tumor cells exhibited significantly decreased patient specificity, but were still highly timepoint specific (**Fig. 2H-I, Fig. S5C**). Tumor cell specific abundance mapping showed TN, MRD, and PD tumor cells occupying distinct regions in the integrated cell embedding **(Fig. 2J)**, but were otherwise intermixed by patient (**Fig. S5A**), suggesting convergent tumor cell states across patients, defined by treatment timepoint.

### Pre-existing poorly differentiated cell populations expand during disease progression

We found that many epithelial cells (both malignant and non-malignant) mapped closely to canonical lung epithelial cell types^19–21^ (**Fig. 3A-C, Fig. S6A)**. These included alveolar type 1 (AT1-like; *AGER^+^CLIC5^+^)* and alveolar type 2 (AT2-like; *SFTPB^+^SFTPC^+^*), basal-like (*TP63^+^KRT17^+^)*, ciliated (*CAPS^+^FOXJ1^+^),* neuroendocrine (*NCAM1^+^NEUROD1^+^*) and cycling cells (*MKI67^+^TOP2A^+^*). Most cell states were present across patients and timepoints (**Fig. S6B-C**), however a population with high expression of hepatocyte markers (hepatocyte-like; *ALB^+^HP^+^*) was derived primarily from liver metastatic samples (**Fig. S6D**); and a cell cluster enriched from four PD patients (PD2, PD12, PD21, PD27) displayed atypical expression from multiple lineages with unknown significance (**Fig. S6A-B**).

**Figure 3.**
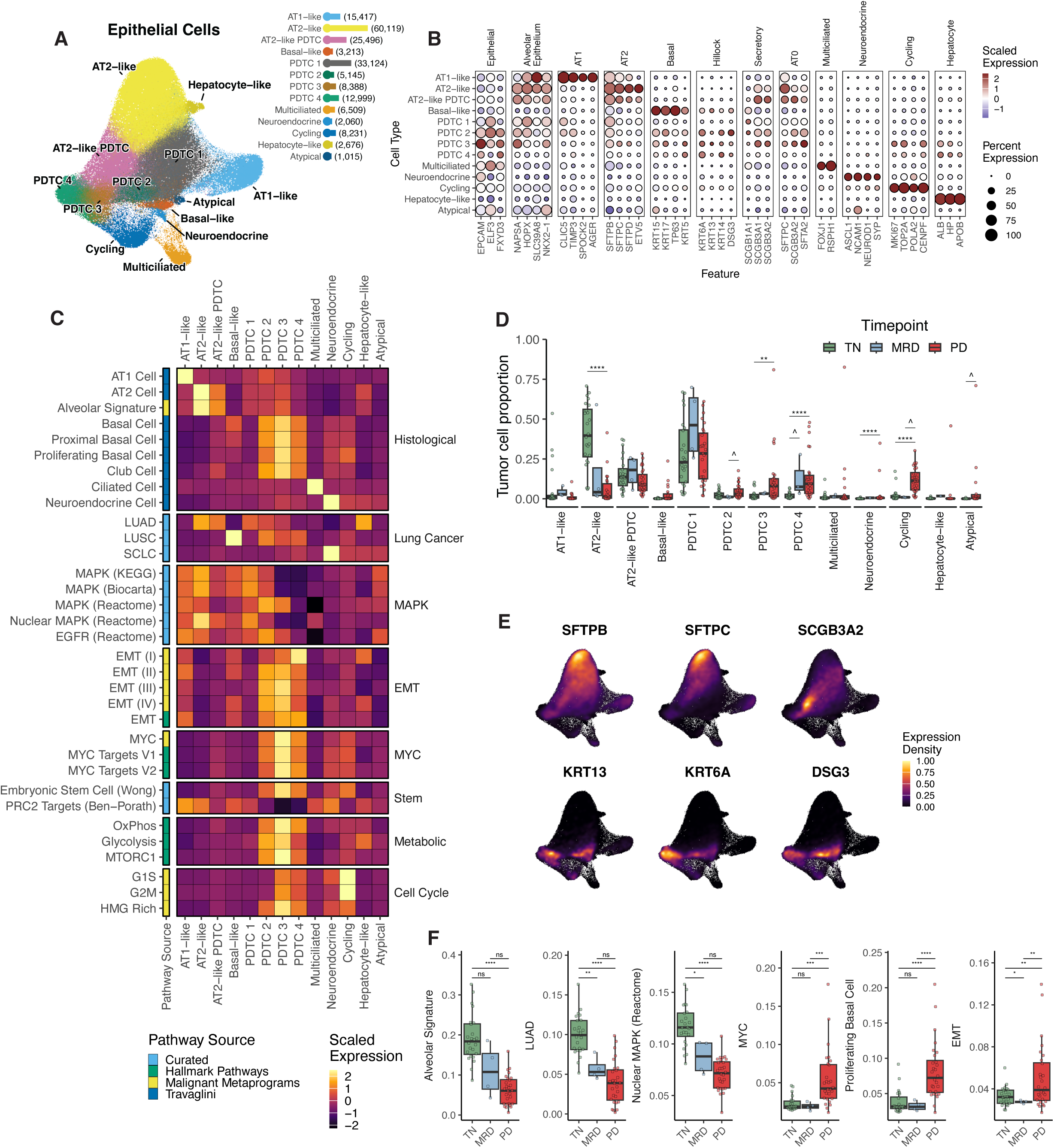
Alveolar cell deprogramming towards stem-like states during disease progression. **A.** UMAP plot of batch corrected malignant and non-malignant epithelial cells, colored by epithelial cell type. Legend barplot indicates the number of each cell type. **B.** Marker dotplots of scaled expression of cell type markers (x-axis) averaged across each epithelial cell type (y-axis). **C.** Heatmap plot of scaled pathway expression across epithelial cell types. The first column indicates the source of each pathway geneset. **D.** Boxplot showing the proportion of tumor cells assigned to each epithelial cell type across treatment timepoints. Each dot represents the proportion of each cell type out of all tumor cells in an individual sample. Wilcoxon rank-sum test significance is encoded as ‘^’: p<0.1, ‘*’: p<0.05, ‘**’: p<0.01 ‘***’: p<0.001, ‘****’: p<0.0001. **E.** Expression density plots for selected cell type markers overlaid on the UMAP from **A**. **F.** Boxplot of signature expression (y-axis) summarized per sample tumor cells (min 20 cells) across treatment timepoints (x-axis). Student’s t-test significance is encoded as ‘ns’: p>=0.05, ‘^’: p<0.1, ‘*’: p<0.05, ‘**’: p<0.01 ‘***’: p<0.001, ‘****’: p<0.0001.

By contrast, we also identified groups of poorly differentiated tumor cells (PDTCs) that were shared across patients and treatment timepoints, but did not closely match canonical lung epithelial cell states. While tumor cells were found across all identified epithelial cell states (**Fig. S6G**), they were particularly enriched in PDTCs, and their relative distribution varied by treatment timepoint (TN, MRD, or PD) (**Fig. 3D**). Most notably, AT2-like tumor cells decreased significantly in PD patients, while PDTCs 2-4, cycling, and neuroendocrine states became more abundant. Interestingly, many of these PDTC states shared properties with recently described stem-like alveolar cell transitional states^22,23^ (SFTPC^low^SFTPB*^+^*SCGB3A2*^+^*) and basal stem-like hillock cells^24,25^ (KRT13*^+^*KRT6A*^+^*DSG3*^+^*; **Fig. 3E**), each implicated in epithelial regeneration after injury in the distal and proximal airways, respectively. PDTC 1, for example, was the most prevalent cell state present in MRD, but was also abundant in TN and PD. These cells expressed markers of both AT1 and AT2 cells (CLIC5*^+^*SFTPB*^+^*), but were distinct from PDTCs 2-4 as they lacked cycling, EMT, MYC and other signatures, while having intermediate MAPK activation. These cells thus may represent the slow-proliferating phenotype characteristic of the drug-tolerant persister state, which are notably present before *and* after treatment^10^.

AT2-like PDTCs, which were also present across timepoints, were characterized by moderately decreased expression of AT2-like markers and increased expression of secretory markers *SCGB3A1* and *SCGB3A2,* compared to AT2-like cells. The remaining four PDTC states (PDTCs 1-4) showed progressive loss of *SFTPB* (**Fig. S6E**), whose expression is associated with distal versus proximal airway localization^23^. Consistent with this, several PDTC states displayed an increase in markers associated with proximal airway cells (basal or hillock cell markers). Specifically, PDTCs 2-4 displayed markedly diminished alveolar and MAPK signatures and exhibited profiles of basal and club cell types, lung squamous signatures, as well as increased EMT, MYC, stemness and metabolic activity (**Fig. 3C**). PDTCs 1 and 2 expressed AT1 markers (CLIC5*^+^*TIMP3*^+^*), whereas PDTC 3 showed relatively higher expression of alveolar transitional cell (AT0) markers and increased cycling signatures. PDTC 4 was the most poorly differentiated PDTC cluster, lacking expression of almost all alveolar epithelial markers.

Aggregating tumor cell measurements per patient, we then quantified differences in tumor cell intrinsic signalling, faceted by treatment timepoint (**Fig. 3F, Fig. S7**). Alveolar, LUAD, and MAPK signatures decreased in a stepwise fashion between TN, MRD and PD patients. In contrast, PD tumor cells were found to have significant increases in MYC, proliferating basal signatures and EMT. Taken together, by defining archetypal tumor cell states across treatment timepoints, we have shown that progression is characterized by loss of alveolar cell identity, and expansion of plastic stem-like tumor cells reminiscent of transitional cell states induced during lung injury. Critically, slower cycling PDTCs exist prior to therapy exposure, expand in MRD, and persist in PD, suggesting that resistant phenotypes may emerge from pre-existing tumor cell plasticity.

### Adenocarcinoma cell identity is linked to MAPK activation

The coincident decrease in alveolar, LUAD, and MAPK signatures accompanying disease progression prompted investigation into how these pathways correlate to one another. This is of particular interest given that tumors resistant to *EGFR* targeted therapies commonly acquire resistance mutations that are thought to sustain MAPK signaling^7^. We first noted that MAPK and LUAD signatures across patients were positively correlated **(Fig. 4A)**. This association was recurrent within patient tumors and was not found with LUSC or SCLC signatures (**Fig. 4B, Fig. S8A**). We validated these findings in an external cohort of single cell sequenced LUAD samples^26^ (n=109), which recapitulated the MAPK and LUAD correlation in both early and advanced stage LUADs (**Fig. 4C**), and in two bulk LUAD tumor sequencing datasets^27,28^ (**Fig. S8B**). Notably, LUAD and MAPK signatures in normal lung AT1 and AT2 cells were anti-correlated, suggesting tumor cell specificity (**Fig. 4D**). Importantly, LUAD, alveolar, LUSC and SCLC gene sets share at most one gene with each other, and with any of the MAPK signatures used, confirming the gene signatures are independent in this regard (**Fig. S8C**).

**Figure 4.**
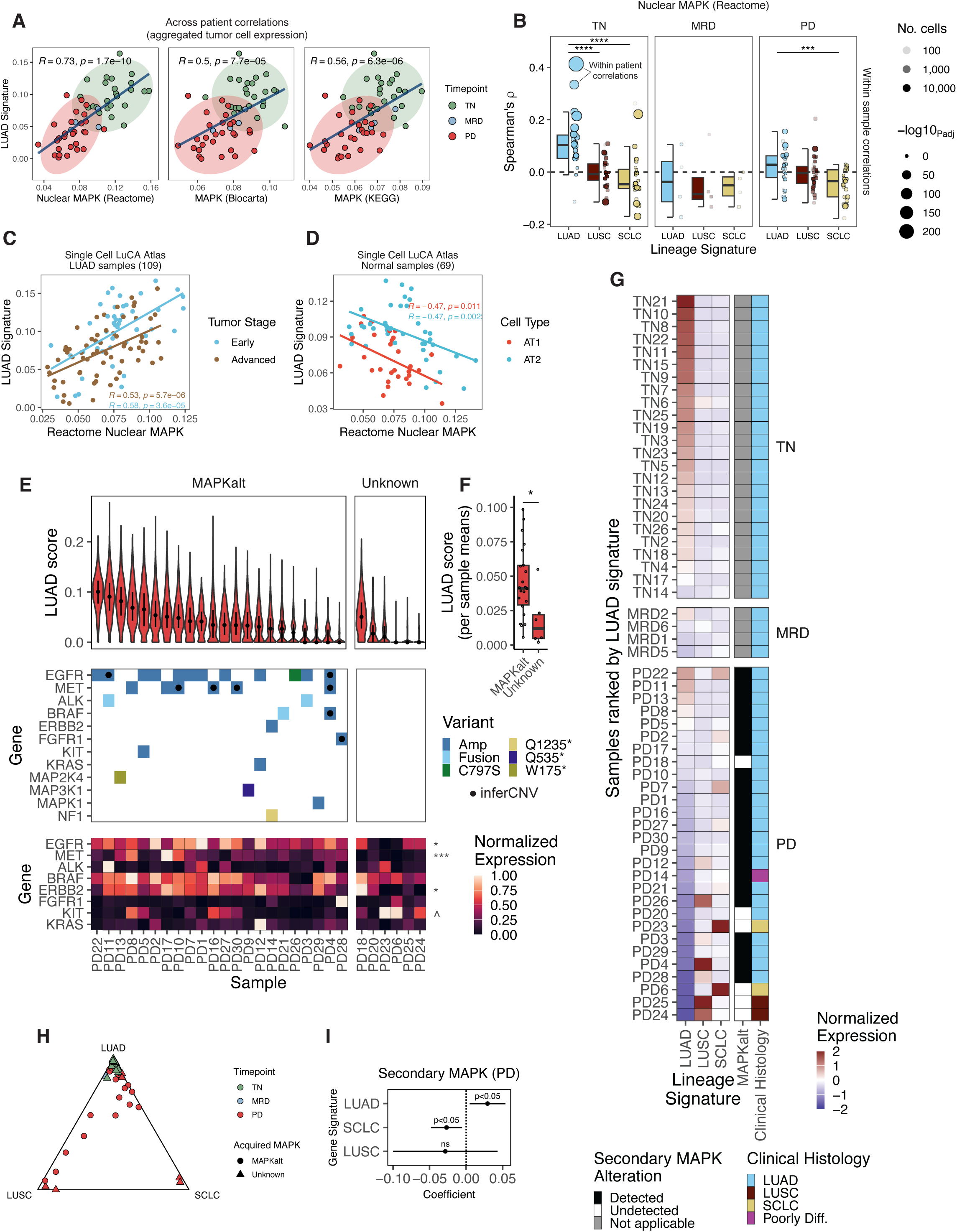
Adenocarcinoma cell identity is linked to MAPK activation. **A.** Scatterplots of MAPK pathway signature expression (3 pathway sources; x-axis) compared to LUAD signature (y-axis) for each patient’s tumor cells (min 20 cells). Points are colored by treatment timepoint, with ellipses surrounding TN and PD distributions. Pearson’s correlation coefficien*t (R)* and significance level are annotated in each panel. **B.** Box and dotplots showing within sample correlations between the nuclear MAPK pathway (Reactome) and LUAD, LUSC and SCLC signatures. Each dot corresponds to a correlation test performed across tumor cells within an individual sample, sized by the level of significance (holm-adjusted), and shaded based on total cell counts. The y-axis indicates Spearman’s ρ. Significance levels of Wilcoxon rank-sum tests comparing the distributions of correlations are encoded as ‘***’: p < 0.001, ‘****’: p < 0.0001. **C-D.** Scatterplots of Reactome Nuclear MAPK pathway expression (x-axis) compared to LUAD signature (y-axis) in tumor and normal cells from an external scRNA dataset^26^. Pearson’s correlation coefficients (*R*) are annotated in each plot, performed separately for early and advanced stage tumors, or normal AT1 and AT2 cells respectively. **E.** Violin plot of LUAD signature scores across PD tumors, faceted by secondary MAPK alterations (dot and linerange indicate median and IQR; top). Oncoprint of MAPK pathway genes and mutation types detected in each PD sample with dots indicating an amplification that was detected by inferCNV (middle). Heatmap of individual MAPK gene (y-axis) expression levels in each PD tumor (bottom). Wilcoxon rank-sum test significance is encoded to the right comparing the MAPKalt versus Unknown as ‘^’: p<0.1, ‘*’: p<0.05, ‘***’: p<0.001. **F.** Boxplot summarizing LUAD signature per sample means between PD tumors with or without detected secondary alterations in the MAPK pathway. Wilcoxon rank-sum test significance is encoded as ‘*’: p<0.05. **G.** Heatmap of LUAD, LUSC and SCLC signature scores in patients tumor cells ordered by decreasing LUAD signature. Detected secondary MAPK alterations for PD patients and clinical histology for patients is annotated on the right. **H.** Ternary diagram showing the relative normalized expression of each histological signature in each sample. **I.** Coefficients and 95% confidence intervals from linear regression modeling the association between secondary MAPK alterations and the expression of each histological signature in PD patients.

We next sought to analyze MAPK signaling in the context of matched clinical panel based genomic sequencing^18^. Secondary alterations were detected in MAPK pathway genes in 22/28 PD patients (n=2 without IMPACT), including amplifications in *EGFR, MET*, *ERBB2, FGFR1, KIT and KRAS, and* fusions in *ALK* and *BRAF* (**Fig. 4E**). Despite the overall decrease in LUAD signature in PD patients, those harboring secondary genomic alterations to the MAPK pathway displayed a significant increase **(Fig. 4F)**. Intriguingly, PD18 had increased LUAD signature expression despite lacking an identifiable MAPK alteration. Further analysis of MAPK pathway genes showed this patient had the highest levels of *ERBB2* signaling among the PD patients (**Fig. 4E**), which might account for the increased LUAD signature measured in this case. In addition to this exception, not all patients with secondary MAPK activating mutations exhibited high LUAD signatures. Examining expression of mutated genes provided clarity in these cases. For example, PD3 had an *EGFR* amplification and *ALK* fusion, but expression of these genes was diminished compared to other cases with these mutations. PD28 harbored an *FGFR1* amplification with high *FGFR1* expression, however amplifications of this gene occur frequently in squamous histologies^29^, suggesting a LUSC-lineage related resistance mechanism. Finally, PD26 harbored a classic gatekeeper *EGFR* C797S on-target resistance mutation, but had significantly diminished expression of other MAPK pathway genes, with the lowest MAPK pathway activity of all PD cases. These results suggest that while genomic MAPK alterations typically correlate with transcriptional activity, there are exceptions in which mutations may no longer be active or where downstream activity may be driven by other mechanisms.

Interrogation of histological signatures revealed a notable enrichment of LUSC and SCLC signatures in PD patients with low LUAD signature, particularly in those patients lacking secondary MAPK alterations (**Fig. 4G-H**). Accordingly, cases clinically defined as histological transformations matched to those with decreased LUAD and high LUSC or SCLC signatures respectively (PD6, PD23, PD24, PD25). We also confirmed the elevated LUSC signature in *FGFR1-*amplified PD28^29^. We modeled the association of MAPK alterations to each signature in PD samples, which revealed a statistically significant association of secondary MAPK alterations with the LUAD signature, and an inverse relationship for SCLC (**Fig. 4I**), consistent with the incompatibility of MAPK signaling with the neuroendocrine phenotype^30^. Overall, detailed analysis of genome-phenome associations has uncovered distinct patterns linking MAPK activity to adenocarcinoma cell identity in both early and advanced stage disease, with sustained activation of MAPK restraining histological lineage plasticity at disease progression.

### Dynamics of lineage plasticity

We next sought to define cell-intrinsic state dynamics of alveolar cell deprogramming through treatment progression. We constructed pseudotime tumor cell trajectories with known histological lineage markers. We defined this manifold as ‘histological pseudotime’ or ‘histotime’ (**Methods**, **Fig. 5A**). The resulting trajectory captured LUAD dedifferentiation, patterned by progressive loss of AT2 lineage markers and decreased MAPK signaling in the truncal branch, along with two distinct branches marked by high expression of squamous and neuroendocrine markers respectively (**Fig. 5B**).

**Figure 5.**
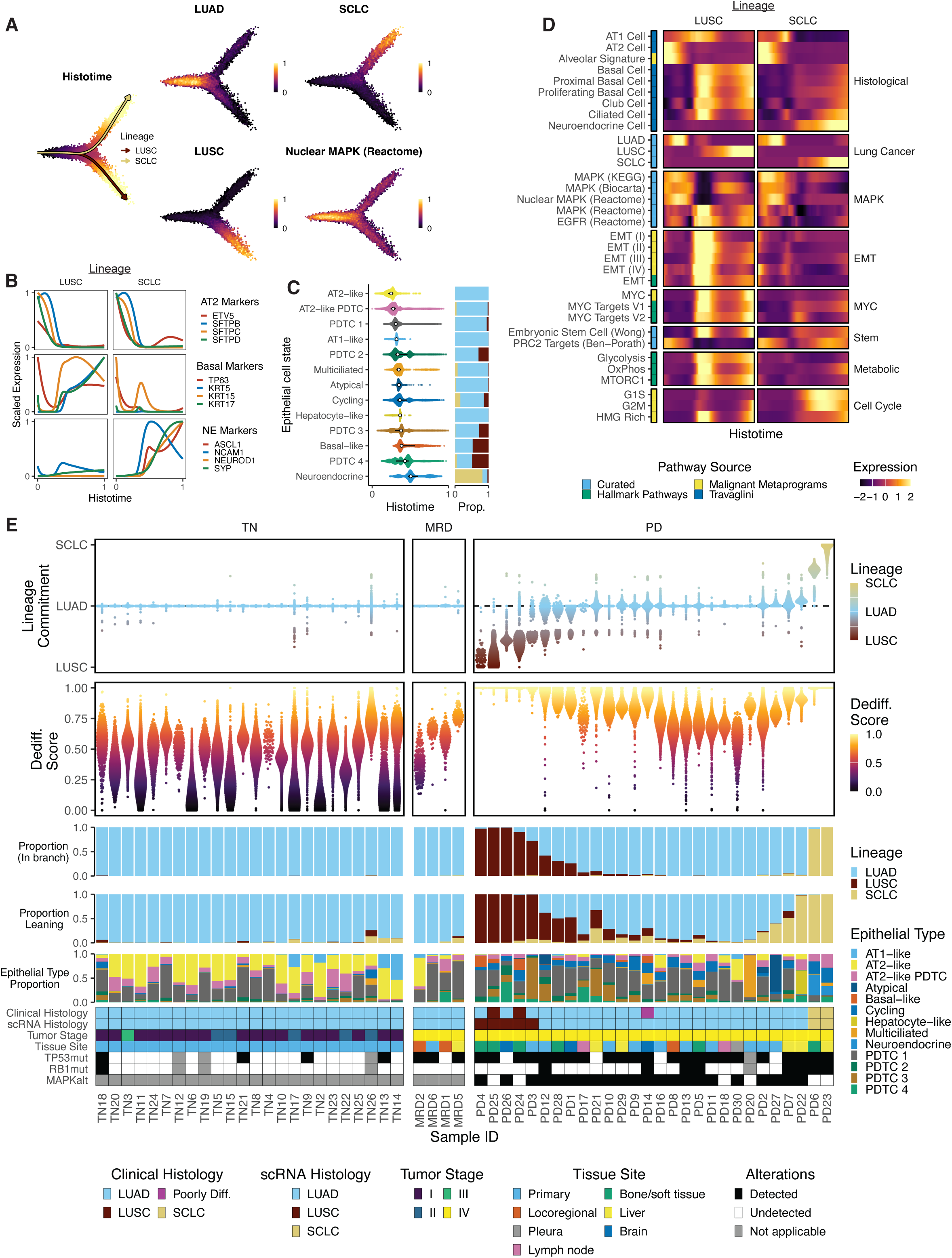
Dynamics of lineage plasticity. **A.** Histotime analysis of malignant cells. Main subpanel (left) colored by histotime with lines indicating the LUSC and SCLC lineage branches. Remaining sub-panels are colored by LUAD, LUSC and SCLC, and Reactome Nuclear MAPK signatures respectively. **B.** Scaled expression of AT2, basal and neuroendocrine markers in the LUSC and SCLC histotime lineage branches respectively. **C.** Violin plot (left) of histotime values across epithelial cell states and proportion (right) of cells for each state belonging to the LUAD, SCLC or LUSC lineage branches resepectively. **D.** Heatmap plot of scaled pathway expression in each histotime lineage (LUSC and SCLC). The first column indicates the source of each pathway geneset. **E.** Violin plot of the relative lineage commitment, LUAD dedifferentiation scores, proportion of cells assigned to, or leaning towards each branch, epithelial cell type proportion, and associated metadata for each patient’s tumor cells.

AT2-like tumor cells were found earliest in histotime, and PDTCs 1-4 were sequentially later (**Fig. 5C**). PDTCs 2-4 had an appreciable proportion of cells in the squamous lineage (28-50%), consistent with their increased expression of stem-like basal markers, while neuroendocrine cells were the latest in histotime, and mapped almost exclusively to the neuroendocrine branch. The two end histological states revealed basal cell, EMT, MYC, and metabolic signatures specifically within the squamous lineage (**Fig. 5D**). Intriguingly, in both LUSC and SCLC lineages, AT1 markers increased as AT2 markers decreased. This shift coincided with increased expression of alveolar transitional cell (AT0) related markers such as *SCGB3A2*, and *SFTA2*, whereas hillock-cell markers were only active within the basal lineage (**Fig. S9A**). Together, these results suggest that emergence of alveolar cell transitional states may generally underpin LUAD cell dedifferentiation and tumor progression, irrespective of histological transformation^31,32^.

We next derived cell-level metrics from histotime, including the degree of squamous or neuroendocrine lineage commitment, a dedifferentiation score representing the degree of alveolar deprogramming in the truncal branch, and the proportion of cells per patient that exhibited features of transformation, but not yet committed to either lineage (‘leaning’, **Methods**, **Fig. 5E**). Lineage plasticity was significantly higher in PD tumors, reflecting a high degree of alveolar dedifferentiation. However, the overall degree of dedifferentiation was lower in MAPK-pathway mutated patients (**Fig. S9B**).

We next classified samples into histological groups based on the predominant lineage. All five cases defined clinically as being squamous transformed (PD24, PD25), small-cell transformed (PD6, PD23), or poorly differentiated (PD14) matched our expression based histological classifications. Beyond these cases, we identified additional PD tumors clinically annotated as LUAD with convincing evidence of lineage plasticity based on histotime, particularly to the LUSC lineage. This included three cases that were predominantly squamous (PD3, PD4, PD26). Notably, PD4 was a suspected squamous transformation, which has now been corroborated by this analysis.

TN tumors were generally more well differentiated compared to PD tumors. However, certain TN patients were found to have rare subsets of tumor cells that displayed far higher dedifferentiation levels, in some cases displaying overt lineage plasticity (e.g. TN9, TN14, TN17, TN26). Given the enrichment of *TP53* mutations in PD (**Fig. S2G**), we wondered whether occurrence of these mutations prior to treatment might prime tumor cell plasticity. Notably, tumor cells from *TP53* mutant patients displayed significantly elevated levels of LUAD dedifferentiation prior to treatment (**Fig. S9C**). These shifts were also observable in the epithelial cell map, with cells from these *TP53* mutant cases showing decreased AT2-like phenotypes, and enrichment for PDTCs 1 and 2 (**Fig. S9D-E**). Taken together, these observations are consistent with a model of modest, pre-existing lineage plasticity in *TP53* mutant tumors and constrained divergence from the adenocarcinoma identity in tumors with acquired MAPK alterations.

## DISCUSSION

Here we report comprehensive single cell profiling of *EGFR*-mutant LUAD before, during, and after treatment with osimertinib, revealing archetypal tumor cell states that define disease progression and treatment resistance. We describe complementary mechanisms of resistance, gated on acquired genomic alterations leading to MAPK pathway activation (**Fig. 6**). While dedifferentiation was common in progressed tumors, MAPK activation appeared to counteract histological plasticity by maintaining LUAD identity. The current paradigm where resistance mechanisms are categorized into on-target (*EGFR* mutated) and off-target (mutations in other receptor tyrosine kinases) overlooks the convergent phenotypic characteristics that accompany MAPK reactivation across these categories – most prominently its association with alveolar cell identity. Critically, this relationship was tumor cell specific and validated using external LUAD datasets, including non-*EGFR* mutant tumors, hinting at broader implications for these findings in other disease contexts.

**Figure 6.**
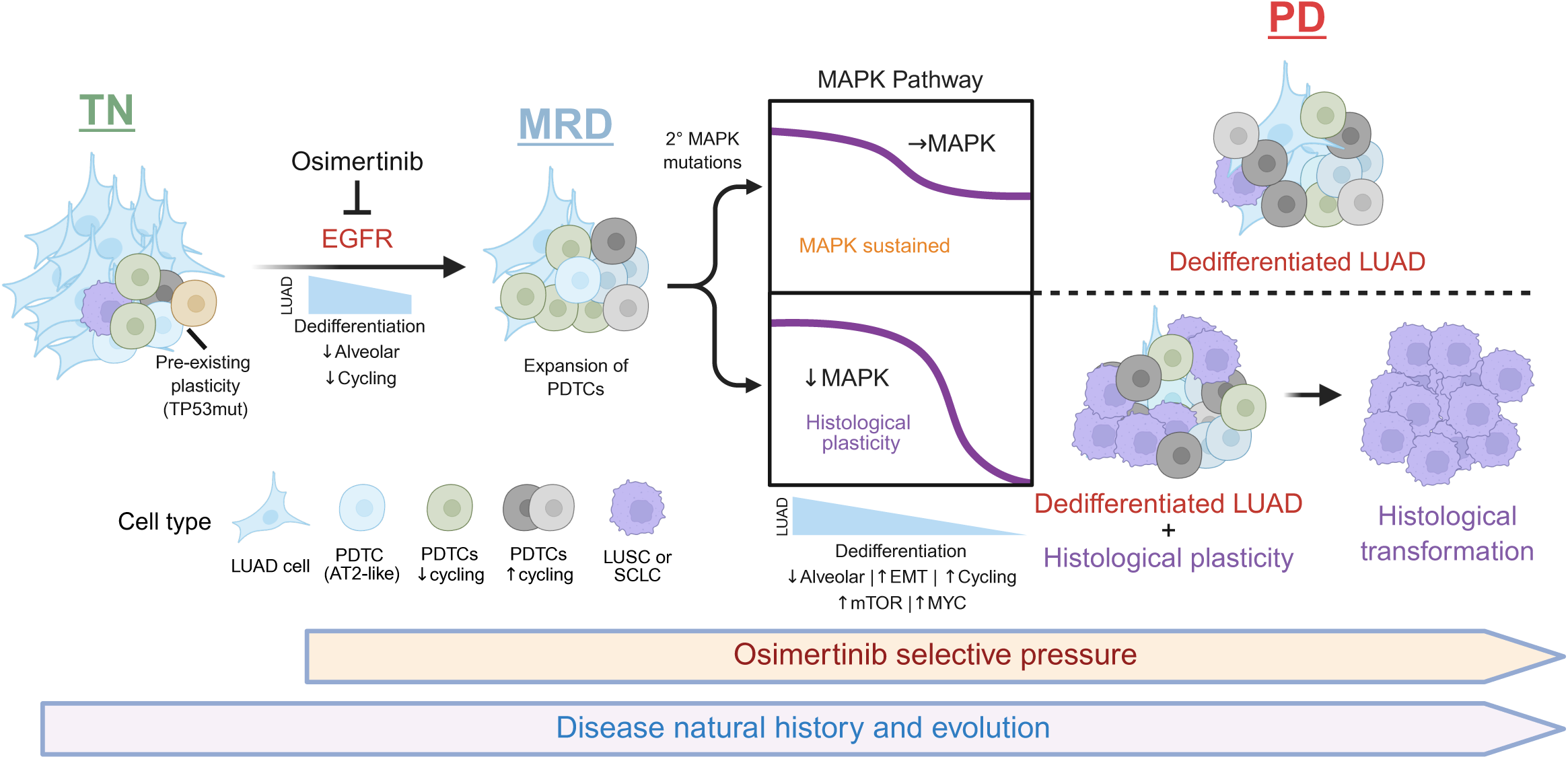
Complementary modes of resistance through MAPK activation and cellular plasticity. Schematic representation of the proposed mechanism of *EGFR* TKI resistance, contingent on MAPK activation. Tumors can harbor pre-existing plastic cells, enriched on *TP53* mutant background, even before therapy exposure. Upon osimertinib treatment, cells begin to lose their alveolar cell identity, and see expansion of slower cycling poorly differentiated tumor cell states in MRD. Upon progression, dedifferentiation is seen broadly regardless of secondary mutations in MAPK pathway genes, along with increased cycling, EMT, mTOR and MYC signatures. Cases with secondary MAPK activation maintain relatively higher LUAD identity. In contrast, cases lacking secondary MAPK activation tend to have higher levels of histological plasticity, in some cases leading to complete histological transformation to LUSC or SCLC. PDTC: poorly differentiated tumor cell.

Granular dissection of tumor cell phenotypes across the treatment timepoints revealed nuanced and progressively dedifferentiated states. We provide additional context for these states, associating them with recently described injury induced stem-like states in the lung, including an alveolar transitional state marked by expression of *SCGB3A2*^22,23^, or basal-like hillock cells marked by *KRT6A*, *KRT13*, and *DSG3* expression^24,25^. These findings support the notion that cellular programs implicated in normal tissue injury repair are co-opted in cancer and can drive disease progression^33^. Notably, the *SCGB3A2*^+^ alveolar transitional cell state has been shown to be inducible *in vitro* by direct *EGFR* inhibition using erlotinib^23^, raising the possibility that *EGFR* targeting therapies may directly induce stem-like survival or transitional states in cancer cells through non-genetic mechanisms. Genetic mechanisms, however, are also implicated in this process, with compelling evidence suggesting that wild-type *TP53* maintains the fidelity of AT2 to AT1 cell transitions during both injury repair and LUAD tumor formation in mouse models^31^. Consistent with this, we observed enrichment of poorly differentiated tumor cells in treatment-naive *TP53* mutant patients, suggesting that *TP53* loss may confer capacity for cellular plasticity – even in disease natural history.

Although our study is limited by unmatched patient samples and the difference in disease staging across treatment timepoints, integration of both canonical and lung tumor specific cell signatures allowed the mapping of tumor cells into a common space that captured disease relevant histological trajectories. This single-cell manifold, which we call ‘Histotime’, captured divergence of tumor cells away from the alveolar lineage towards neuroendocrine and squamous lineages. Notably, Histotime, corroborated all clinically defined cases of histological transformation, and also nominated additional cases with convincing evidence of transformation. We also found subsets of cells with evidence of histological plasticity in several PD tumors that were designated adenocarcinoma on clinical review, suggesting that histological plasticity may be more prevalent than anticipated. Whether these populations are the result of residual tumor subclones lacking secondarily acquired MAPK alterations, new tumor subclones arising from ongoing clonal evolution, or stochastic transcriptional heterogeneity inherent to progressive disease remains an open question.

Poorly differentiated cells, which harbored markers of stemness, EMT, MYC, mTOR and altered metabolism, could be identified before treatment, albeit at lower frequency. These observations suggest that *EGFR*-mutant LUADs may have some degree of inherent plasticity, consistent with some treatment-naïve tumors presenting with mixed histology^34,35^. Indeed, histotime analysis revealed rare cell subsets in TN tumors that mapped to neuroendocrine or squamous histologies. Variation in transcriptional phenotypes prior to treatment, even among clonally related cells, could act as a substrate for differential therapeutic selection^36,37^. This could occur through intrinsically resistant cell states that are selected, or through adaptation to the therapeutic stimulus facilitated by pre-existing plasticity. Some combination of intrinsic cell plasticity, coincident with the timing of additional genetic alterations, likely governs the exact mode of resistance on a per case basis.

In summary, we propose a model where osimertinib induces the expansion of poorly differentiated, stem-like cell states with suppressed MAPK signaling and with the ability to persist on treatment. Such states exhibit potential to drive tumor resistance by either (1) MAPK signaling reactivation and maintenance of the LUAD phenotype or (2) transitions to an alternative histologic phenotype. The described interplay of plasticity, intratumoral heterogeneity and MAPK signaling provides a deeper understanding of how lung adenocarcinomas become resistant to targeted therapy, and should inform future therapeutic approaches tailored to prevent or delay patient relapse.

## METHODS

### Sample collection and pre-processing

All samples were collected under IRB approved protocols including #12-245 and 06-107. We queried for available *EGFR*-mutant lung cancer samples that were treatment-naive, obtained on-treatment on osimertinib or collected at progression on osimertinib. The range of dates of sample collection were between 12/8/2017 and 3/9/2022. 5/26 pre-treatment (TN) and 5/6 on-treatment (MRD) patients were treated with osimertinib and eventually progressed. All 30 PD samples were collected after treatment with osimertinib at the time of radiographic progression. Tumor tissue samples were collected at the time of biopsy or surgical resection and were flash-frozen. The complete cohort oncoprint depicting all detected mutations is provided in **Fig. S2A** and summarized in **Table S2**.

### Sample processing and snRNA Library Preparation

Frozen tumor samples (∼50-200 mg) were used for nuclei isolation. Nuclei extraction was performed following the protocol by Masilionis et al. (https://www.protocols.io/view/nuclei-extraction-for-single-cell-rnaseq-from-froz-b7vxrn7n) using the Singulator 200 (S2 Genomics). Extracted nuclei were stained with 7-aminoactinomycin D (Invitrogen) and FACS-sorted for further purification. snRNA-seq was then performed on FACS-sorted nuclei suspensions using the Chromium instrument (10X Genomics), following the user manual for the 3’ v3.1 Reagent Kit. Each sample, containing approximately 10,000 nuclei at a final dilution of ∼1,000 cells/µl, was loaded onto the cartridge according to the manufacturer’s instructions. During the reverse transcription (RT) step, individual transcriptomes of encapsulated cells were barcoded, and the resulting cDNA was purified using DynaBeads, followed by amplification per the protocol. The PCR-amplified product was then fragmented, A-tailed, purified with 1.2X SPRI beads, ligated to sequencing adapters, and indexed via PCR. The indexed DNA libraries underwent double-size selection purification (0.6–0.8X) using SPRI beads before sequencing on the Illumina NovaSeq S4 platform. Sequencing parameters included R1 – 26 cycles, i7 – 8 cycles, and R2 – 70 cycles or higher, with an average sequencing depth of approximately 200 million reads per sample.

### snRNA sequence alignment and gene counting

Raw 10X sequencing data were aligned using Cellranger (version 7.1.0), which also performed barcode filtering and unique molecular identifier (UMI) gene counting using the 10X GRCh38 reference transcriptome.

### snRNA Quality control

Filtered barcode matrices were first screened for doublets using the scDblFinder R package (version 1.16)^38^ using default settings. Counts were then merged and loaded using Seurat (version 5), and filtered for high quality cells using the following parameters:

- Minimum read count per cell: 1000
- Minimum genes detected per cell: 500
- Maximum genes detected per cell: 7,500
- Maximum mitochondrial read percentage per cell: 20%
- Remove called doublets by scDblFinder

Sample level quality control was then performed using the following parameters:

- Remove any samples with fewer than 300 high quality cells
- Remove any samples with >70% sequencing saturation that failed any one of:

◦ Mean reads per cell less than 25,000
◦ Median genes per cell less than 900
◦ Median UMI counts per cell less than 1000

In total, five of the initially sequenced 67 samples were removed due to quality concerns, leaving a total of 385,770 high quality cells from 62 samples for downstream analysis.

### snRNA data processing and batch correction

Read count log normalization, variable feature selection, count scaling, and PCA were performed using Seurat’s default parameters. Seurat’s `FindNeighbors` and `RunUMAP` were run using the first 30 principal components to generate a non batch corrected cell embedding. Batch integration was performed across patients using Seurat’s sketch integration workflow, sketching 3,000 cells per sample and using the reciprocal principal component analysis (RPCA) integration method. Seurat’s `FindNeighbors` and `RunUMAP` were run using the first 30 RPCA components to generate a batch corrected cell embedding. Cohort-wide clustering was performed using louvain clustering at a resolution of 0.25 on the shared nearest neighbor graph generated using the first 30 RPCA dimensions. Cell cycle scores were computed using the Seurat `CellCycleScoring` function.

### Cell type annotation

Cohort-wide cluster specific markers were found using Seurat’s `FindAllMarkers` function. One cluster was found to contain cycling cells from different cell compartments, and was therefore sub-clustered to yield cycling epithelial, myeloid, and lymphoid cells. Clusters were manually reviewed and grouped into major cell type compartments: (1) epithelial cells; (2) T and NK cells; (3) myeloid cells; (4) fibroblasts; (5) endothelial cells (6) B and plasma cells. Rare cell types specific to metastatic sites, such as neuronal cells, liver hepatocytes were also found, and grouped into the ‘Other’ category, and excluded from downstream analysis.

Epithelial cells were subclustered using Louvain clustering at a resolution of 0.5 using the batch integrated neighbor graph. Clusters were then manually annotated, with non-distinct cell types being merged according to marker expression.

### Identification of malignant cells

Malignant cells were identified on the basis of copy number alterations using `inferCNV` (version 1.14)^39^. We created a reference cell set by equally sampling coarse cell types from all other samples (5 per sample across major cell compartments: epithelial, stromal, immune and endothelial). InferCNV smoothed and denoised counts were then aggregated to 10Mb bins, reduced to 50 dimensions by PCA, and then clustered using leiden clustering at 0.3 resolution on the SNN graph. Each sample was then manually inspected to identify malignant cells, determined by presence of copy number variation, patient specificity of CNV signal, near complete enrichment of epithelial cells in clusters, and when available, guided by bulk DNA sequencing derived copy number signal. Six samples (TN1, TN16, MRD3, MRD4, PD15, PD19) had no identifiable tumor cell population using this approach.

### Cell type specificity analysis

Patient and timepoint specificity scores were calculated using either batch uncorrected or batch corrected SNN graphs by calculating the fraction of each cell’s nearest neighbor, divided by the expected fraction in the patient or timepoint subgraph. Subsampling was inversely weighted by the size of each identity (i.e. sample id and cell type or timepoint and cell type) to ensure equal representation in the subgraph, even for lower frequency cell types or samples. For each cell, the sum of the weights of neighboring cells that match the desired identity (for example, tumor cells neighboring other tumor cells from the same patient or timepoint) is divided by the total weights of all neighboring cells to produce a normalized per cell specificity score.

### Geneset signature scoring

Genesets were curated from the molecular signatures database^40^ using the R package `msigdbr` (v2023.1.1), malignant metaprograms^41^, and manually from prior studies of lung cancer^42^. We generated a custom SCLC signature using Cancer Cell Line Encyclopedia expression data ^43^ by subsetting for LUAD, LUSC and SCLC cell lines, running Seurat’s `FindAllMarkers` function and then filtering for genes with a percent difference in expression of at least 10%, that were expressed in at least 50% of SCLC cell lines and less than 50% of LUAD and LUSC cell lines, with an adjusted p-value of less than 0.05. This procedure yielded 38 genes highly specific to SCLC and are provided in **Table S3**. Genesets were scored using `UCell` (v2.10.1)^44^ using default parameters.

### Expression density plots

Expression density plots were created using Nebulosa (v1.0.1)^45^ to visualize expression of select markers as a 2d kernel density estimation over cells.

### Lung cancer atlas data analysis

Lung cancer cell atlas data was downloaded^26^ and signatures scored identically to our snRNA dataset. We filtered for cells annotated as malignant from lung adenocarcinoma samples, or AT1 and AT2 cells derived from normal tissue samples prior to pseudobulking mean expression values for each signature for each sample, and computing correlations.

### MSK-IMPACT targeted DNA sequencing

Genomic DNA from tumor tissue and matched normal blood (n=57) underwent hybrid capture and sequencing to a median coverage of 575x (range 124-954). Variant calling and copy number analysis were performed according to the MSK-IMPACT clinical pipeline (https://github.com/mskcc/Innovation-IMPACT-Pipeline). FACETS^46^ was used in a subset of samples (n=47) to estimate tumor ploidy, fraction genome altered, fraction of genome with loss-of-heterozygosity (LOH), and whole genome doubling (WGD). Enrichment of *TP53* and *RB1* mutations in PD versus TN samples was measured using Fisher’s test.

### MAPK alteration analysis

MAPK altered samples were identified on the basis of IMPACT DNA sequencing, filtered for oncoKB annotated variants known to drive MAPK pathway expression and resistance to *EGFR* inhibition included in **Fig. 4E**. In certain cases where the IMPACT DNA sample was taken prior to osimertinib exposure, amplifications detected from inferCNV on snRNA data were used.

### Histotime analysis

To generate histologically relevant tumor cell trajectories, we first restricted the input feature space to genes related to normal and cancer related lineage markers. Feature selection involved utilizing existing genesets, including the LUAD, LUSC and SCLC signature sets, the alveolar cell malignant metaprogram geneset^41^, and marker genes from the Human Lung Cell Atlas^20^. The final list included 179 genes (**Table S4**). Cells with fewer than 20 total counts across these genes were removed, leaving 127,969 tumor cells for downstream analysis. We applied PCA with a rank of 10, whereupon examination of the first two PCs revealed an association with the tumor histological signatures. We applied gaussian mixture model based clustering using `Mclust` (v6.1)^47^ with default parameters, merging down to four clusters prior to performing trajectory inference using `Slingshot` (v2.10)^48^. To find gene expression patterns along the histotime gradient, we used `TradeSeq` (v1.16.0)^49^ to fit a negative binomial generalized additive model over genes, subsampling to 2,000 cells for each cluster from `Mclust`, and only including genes with at least 1 count in at least 50 cells. To measure pathway level expression over histotime, we applied `UCell` (v2.10.1)^44^ over smoothed gene expression values emitted from `TradeSeq`. Lineage commitment was measured for each cell by calculating the difference in lineage weights for each cell and multiplying by histotime. Dedifferentiation scores were calculated by subsetting for cells assigned to the truncal LUAD branch, clipping the bottom 5% and top 1% of histotime values, and renormalizing the values to a range of 0-1. snRNA based histological classifications were assigned based on the most abundant branch assignment for each sample.

### Code availability

All code required to reproduce the analyses and figures in this paper are available on GitHub (https://github.com/shahcompbio/egfr_nucseq).

## ACKNOWLEDGEMENTS

This study was supported by NIH T32 CA1600001 (to AQV), NCI R01 CA264078 (to HAY/CMR), NCI R35 CA263816 (to CMR), NCI U24 CA213274 (to CMR), by the American Lung Association (to AQV), by Druckenmiller Center for Lung Cancer Research (AQV, and CMR), and the Halvorsen Center for Computational Oncology (MZ, and SPS). We acknowledge the use of the PPBC Biobank, Pathology Core Facility, and Integrated Genomics Operation Core, funded by the NCI Cancer Center Support Grant (P30 CA008748), Cycle for Survival, and the Marie-Josée and Henry R. Kravis Center for Molecular Oncology. JJL is supported by NIH training grant T32CA009207. SS is supported by NIH grant 5K99CA277562-02.

## AUTHOR CONTRIBUTIONS

Conceptualization: AQV, HAY. Data curation: MZ, AQV, AP, CF, HAY. Formal Analysis: MZ, SS. Funding acquisition: CMR, AQV, HAY. Investigation: MZ, AQV, SS, NC, JJL, IM, OC, PM, RC, HAY. Project administration: AQV, AP, CF, HAY. Resources: AQV, AMP, CMR, SPS, HAY. Software: MZ, SS, NC. Supervision: AQV, AMP, CMR, SPS, HAY. Visualization: MZ, AQV. Writing – original draft: MZ, AQV, SPS, HAY. Writing – review & editing: MZ, AQV, NR, CMR, SPS, HAY.

---

**Supplementary Figure 1.**
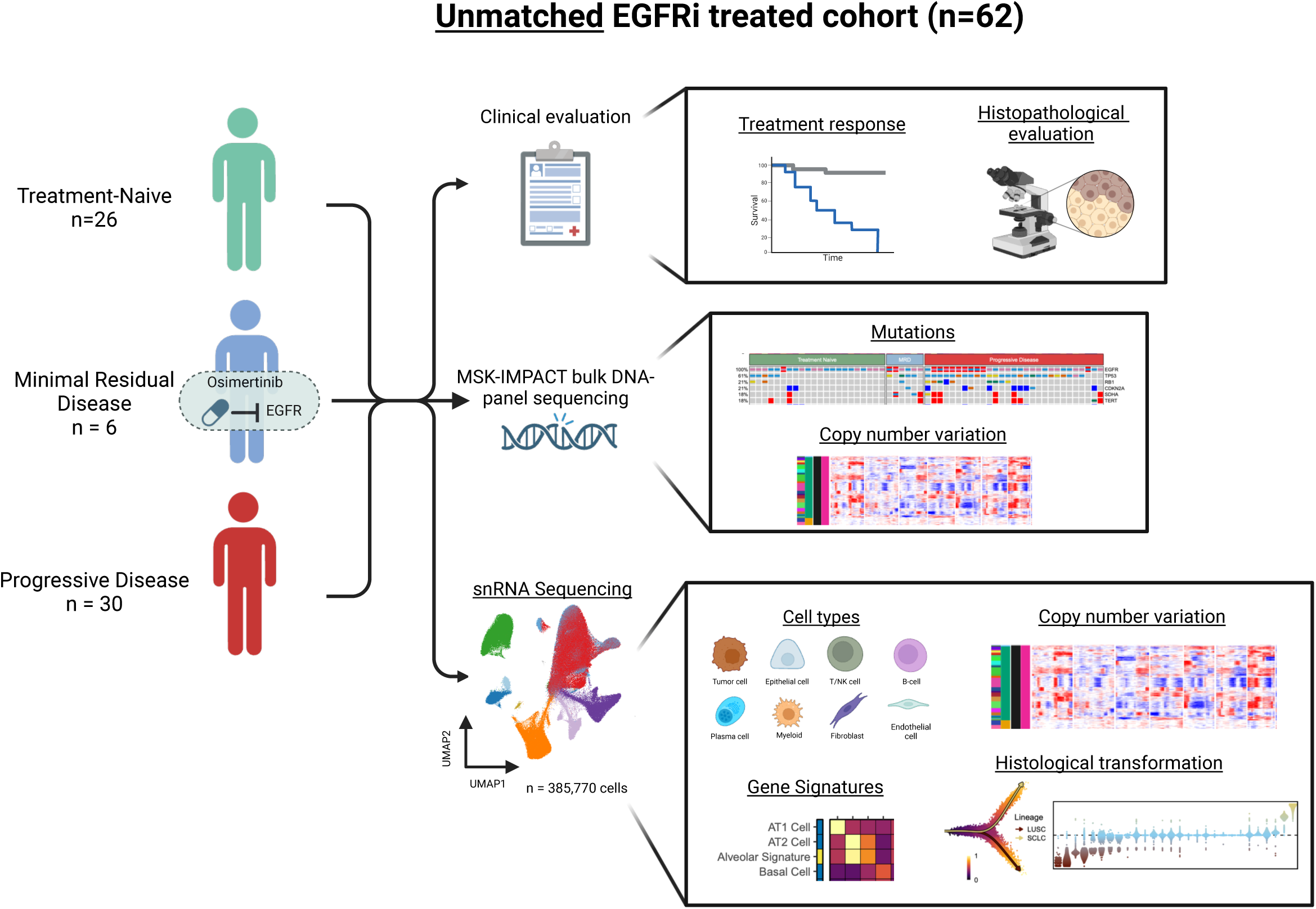
Study diagram. Diagram depicting cohort overview, molecular assays, and analytical approaches used in our study.

**Supplementary Figure 2.**
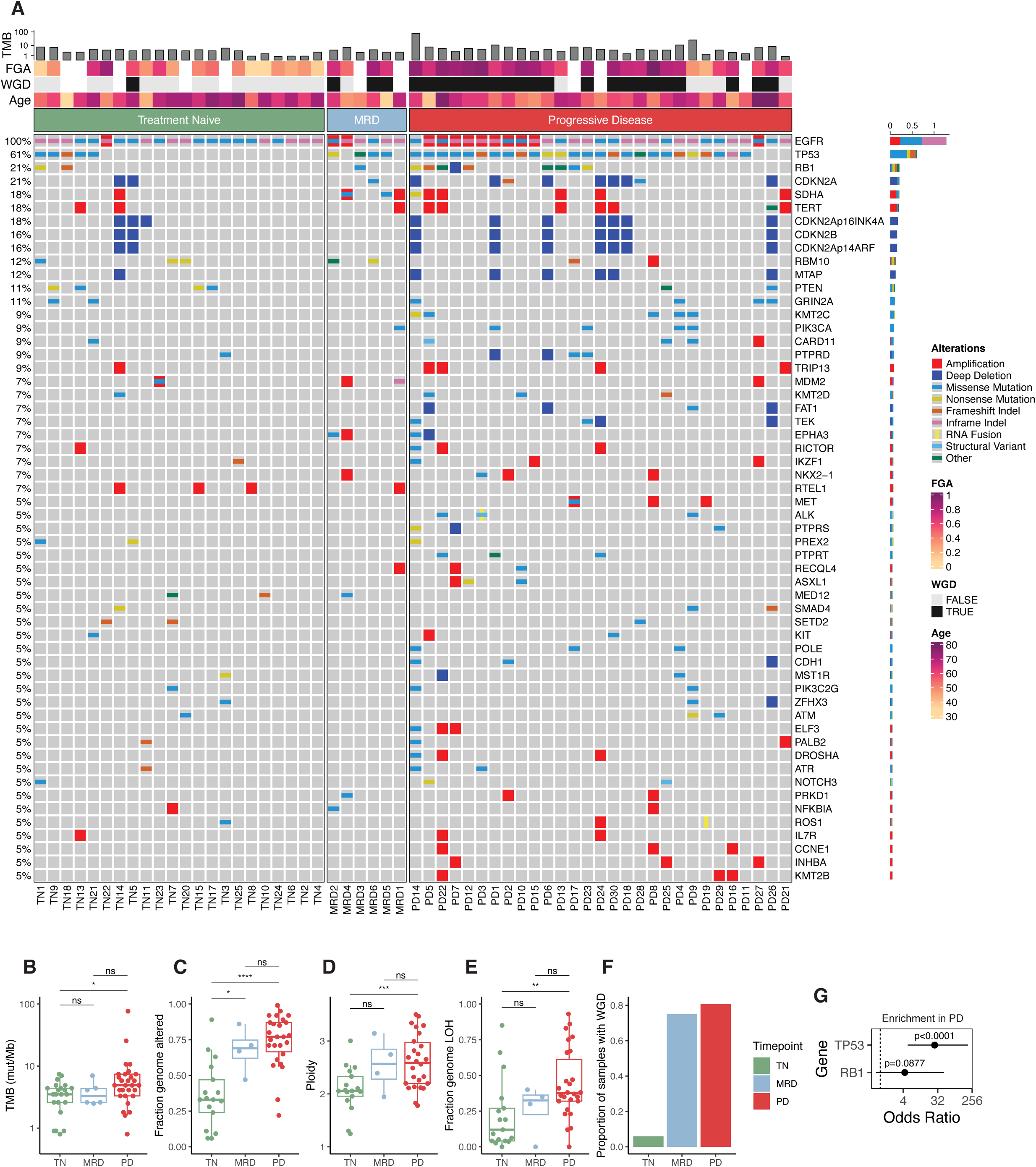
Tumor mutation profiles. **A.** Cohort oncoprint showing mutations detected in each sample, colored by mutation type. Tumor mutation burden (TMB), fraction genome altered (FGA), whole genome doubling (WGD) status, and age are summarized in the top margin of the plot. Per gene mutation frequency is summarized in the right margin. Only mutations present in at least 5% of samples are included. The complete list of mutations are included in **Table S2**. **B-E**. Boxplots of TMB **(B)**, FGA **(C)**, ploidy **(D)**, and loss-of-heterozygosity (LOH) fraction **(E)** summarized across treatment timepoints. Wilcoxon’s rank-sum test significance is encoded as ‘ns’: p>=0.05, ‘*’: p<0.05, ‘**’: p<0.01 ‘***’: p<0.001, ‘****’: p<0.0001. **F.** Barplot of fraction of samples with WGD across treatment timepoints. **G.** Log-odds ratio of enrichment of *TP53* and *RB1* alterations in PD samples.

**Supplementary Figure 3.**
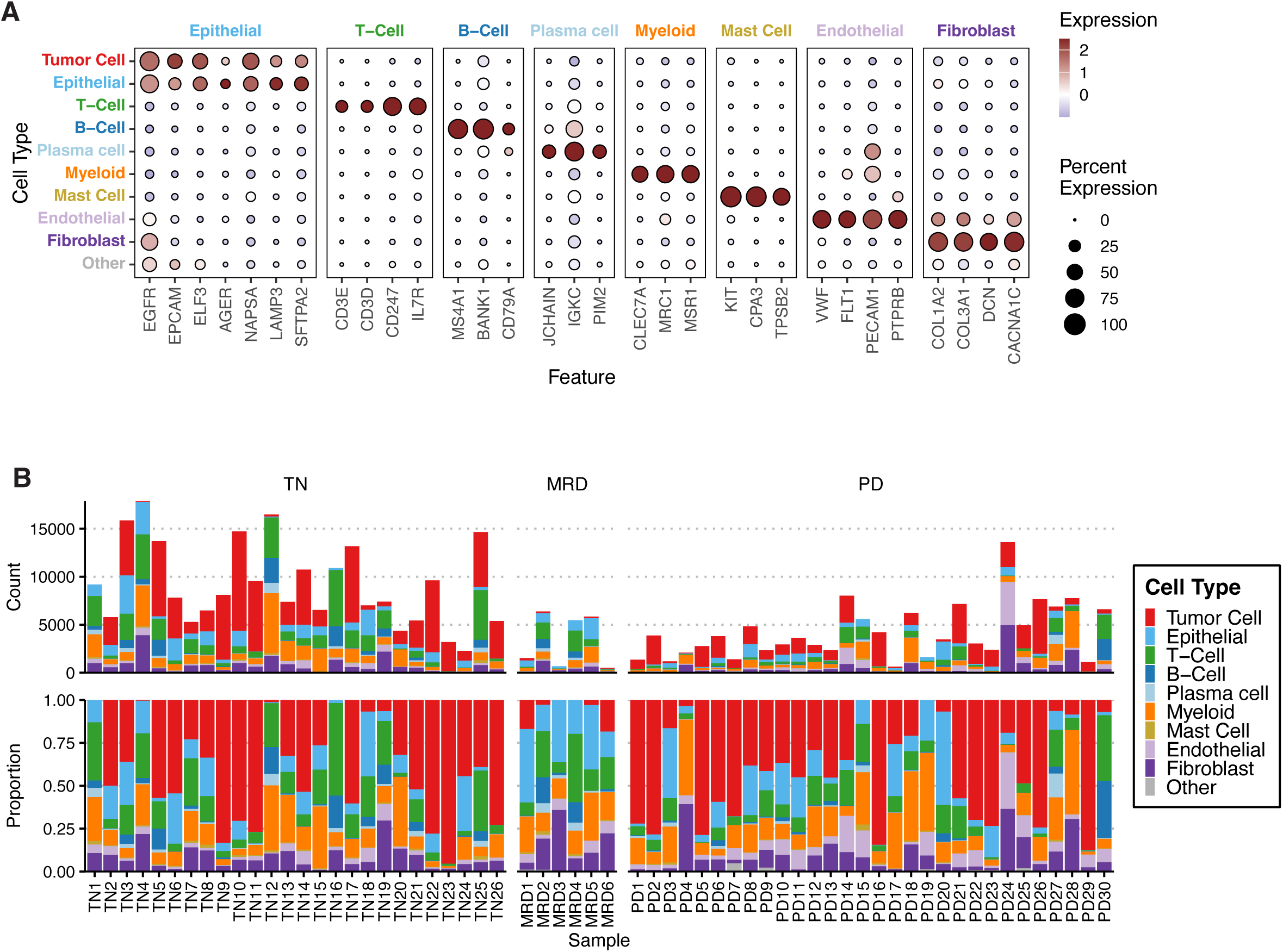
Cohort cell type composition. **A.** Marker dotplots of scaled expression of cell type markers (x-axis) averaged across each cell type compartment (y-axis). **B.** Barplot of major cell type counts (top) and proportions (bottom) in each patient.

**Supplementary Figure 4.**
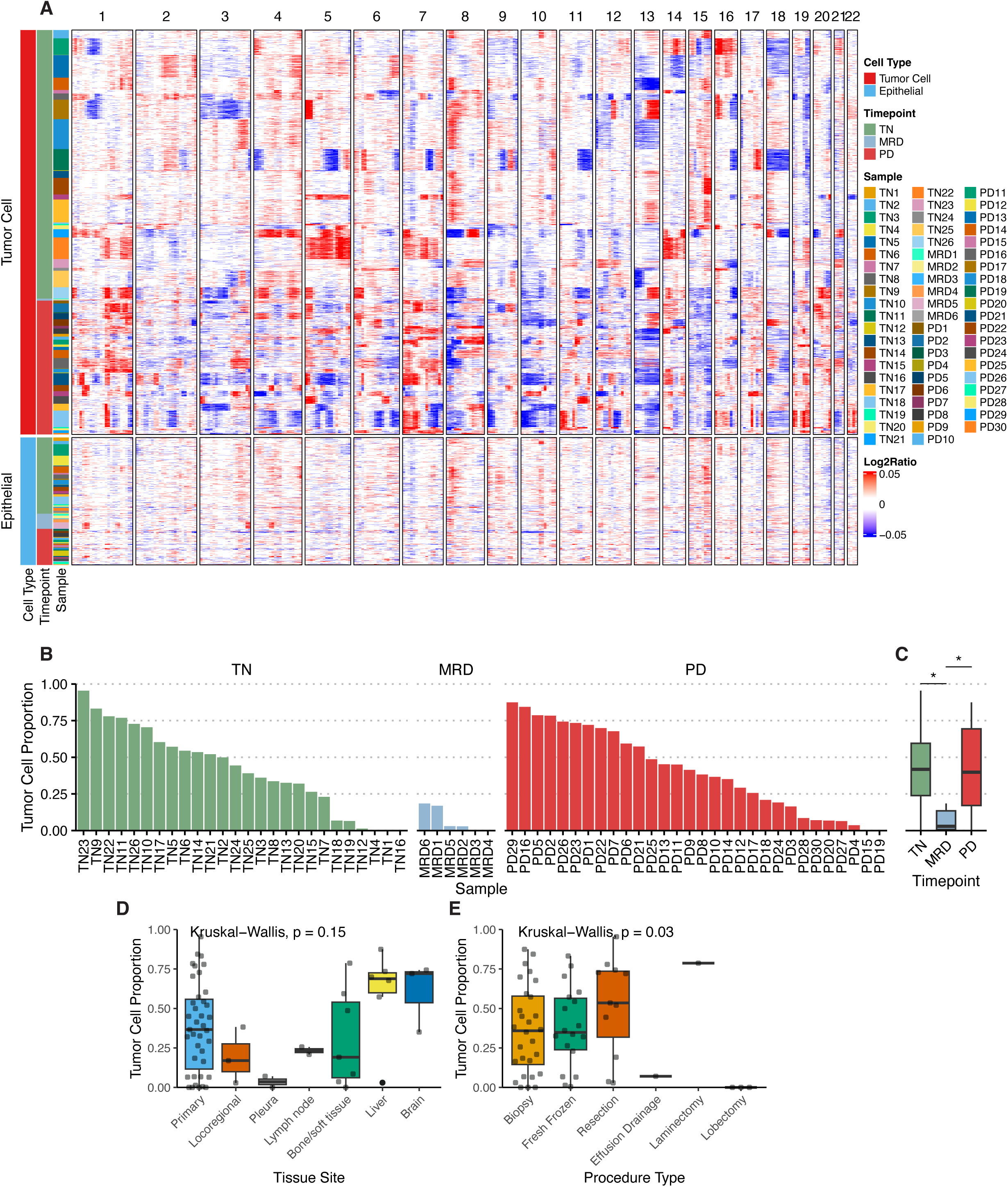
Malignant cell identification. **A.** Heatmap of per cell copy number inferred from snRNA data for malignant and non-malignant epithelial cells. Each row represents an individual cell, and columns are copy number segments across the chromosomes. Cell type, treatment timepoint and samples are annotated in the left margin. **B.** Barplot of tumor cell proportion for each patient (out of all cells). **C.** Boxplot summaries of tumor cell proportion for each treatment timepoint. Wilcoxon rank-sum test significance is encoded as ‘*’: p<0.05. **D-E.** Boxplots of tumor cellularity grouped by tissue site **(D)** and procedure type **(E)**.

**Supplementary Figure 5.**
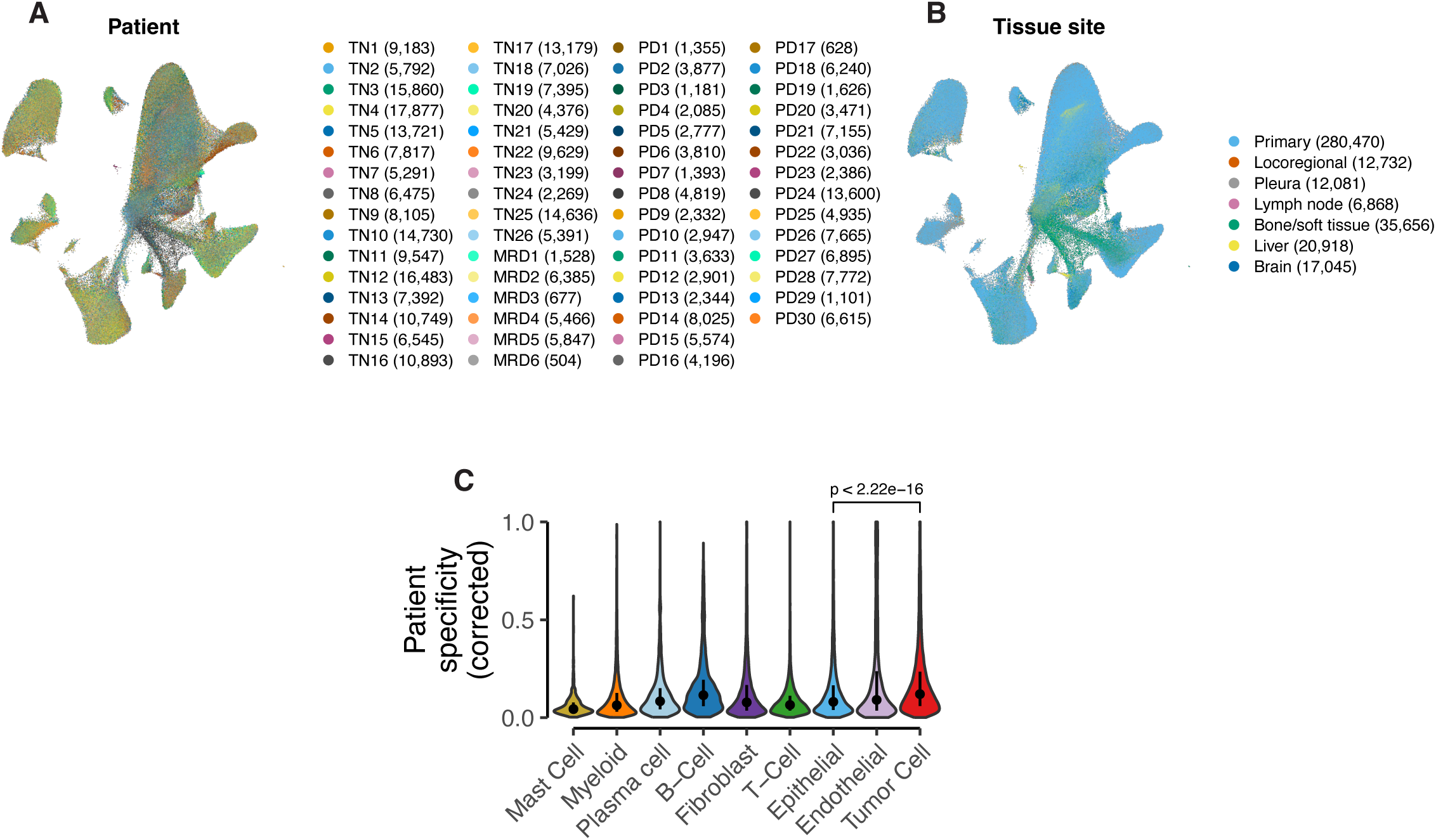
snRNA data integration A-B. UMAP plot of cells after batch correction, colored by patient **(A)** and tissue site **(B)**. **C.** Patient specificity scores for each cell type on batch corrected data.

**Supplementary Figure 6.**
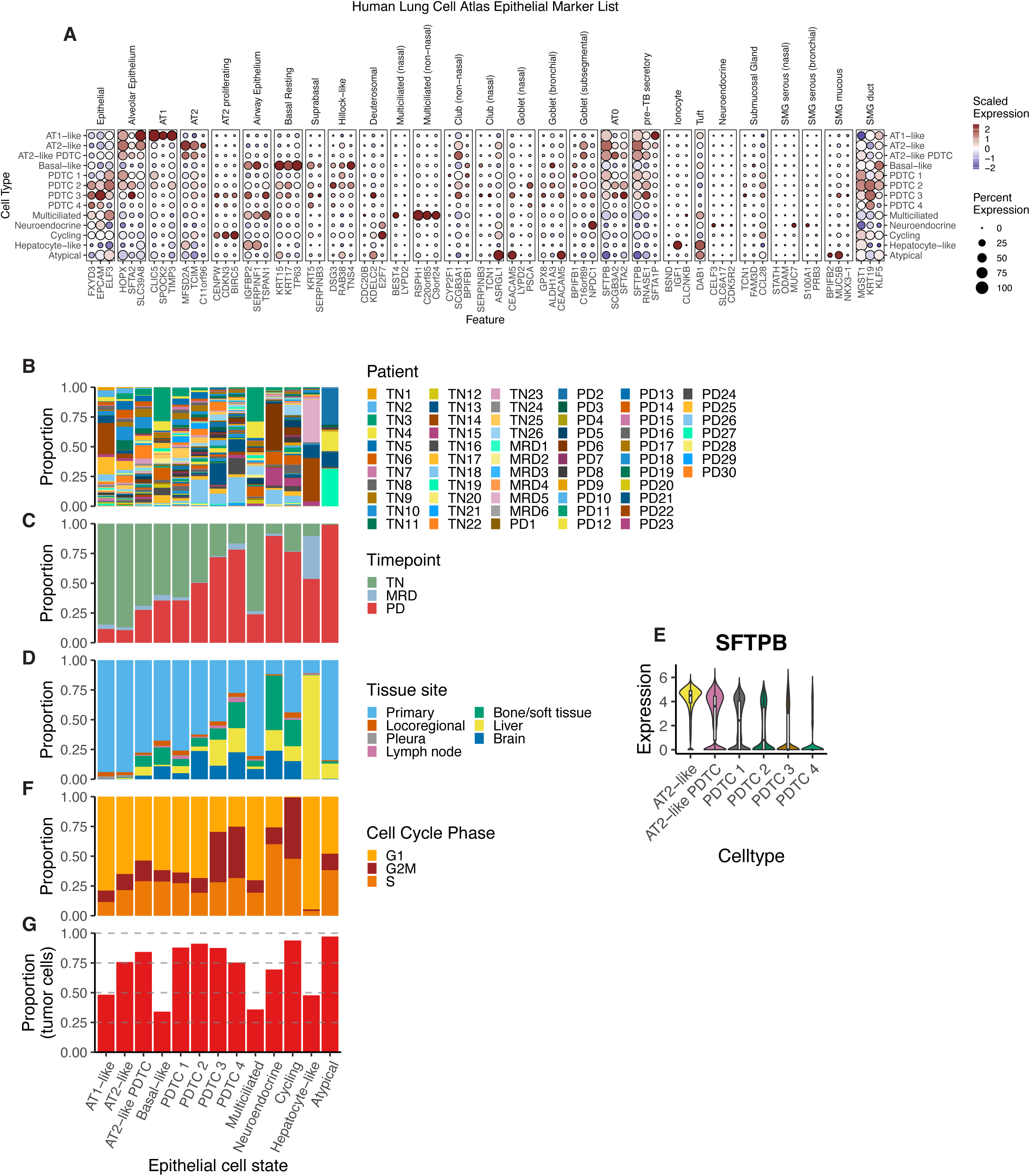
Epithelial cell states. **A.** Marker dotplots of scaled expression of cell type markers (x-axis) averaged across each cell type compartment (y-axis). Markers are derived from the Human Lung Cell Atlas^20^. **B-D.** Barplots showing the proportion of each patient **(B)**, timepoint **(C)** and tissue site **(D)** for each epithelial cell state. **E.** Violin plot of *SFTPB* expression levels across selected epithelial cell states. **F.** Barplot showing the proportion of cells in each cell cycle phase for each epithelial cell state. **G.** Barplot showing the proportion of tumor cells in each epithelial cell state.

**Supplementary Figure 7.**
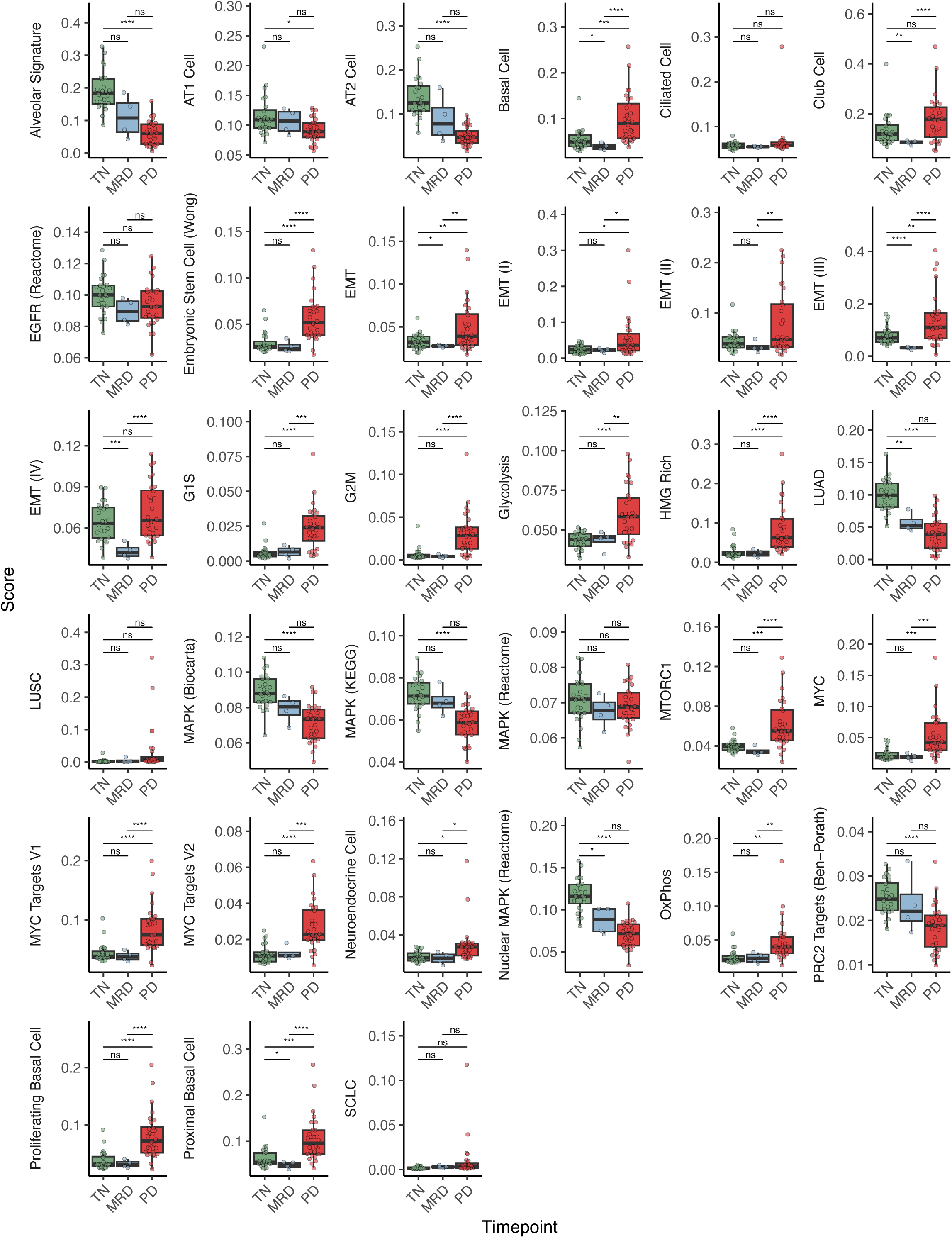
Tumor cell intrinsic signatures. Boxplot of signature expression (y-axis) summarized per patient tumor cells (min 20 cells) across treatment timepoints (x-axis) for the signatures highlighted in Fig. 3C. Student’s t-test significance is encoded as ‘ns’: p>=0.05, ‘^’: p<0.1, ‘*’: p<0.05, ‘**’: p<0.01 ‘***’: p<0.001, ‘****’: p<0.0001.

**Supplementary Figure 8.**
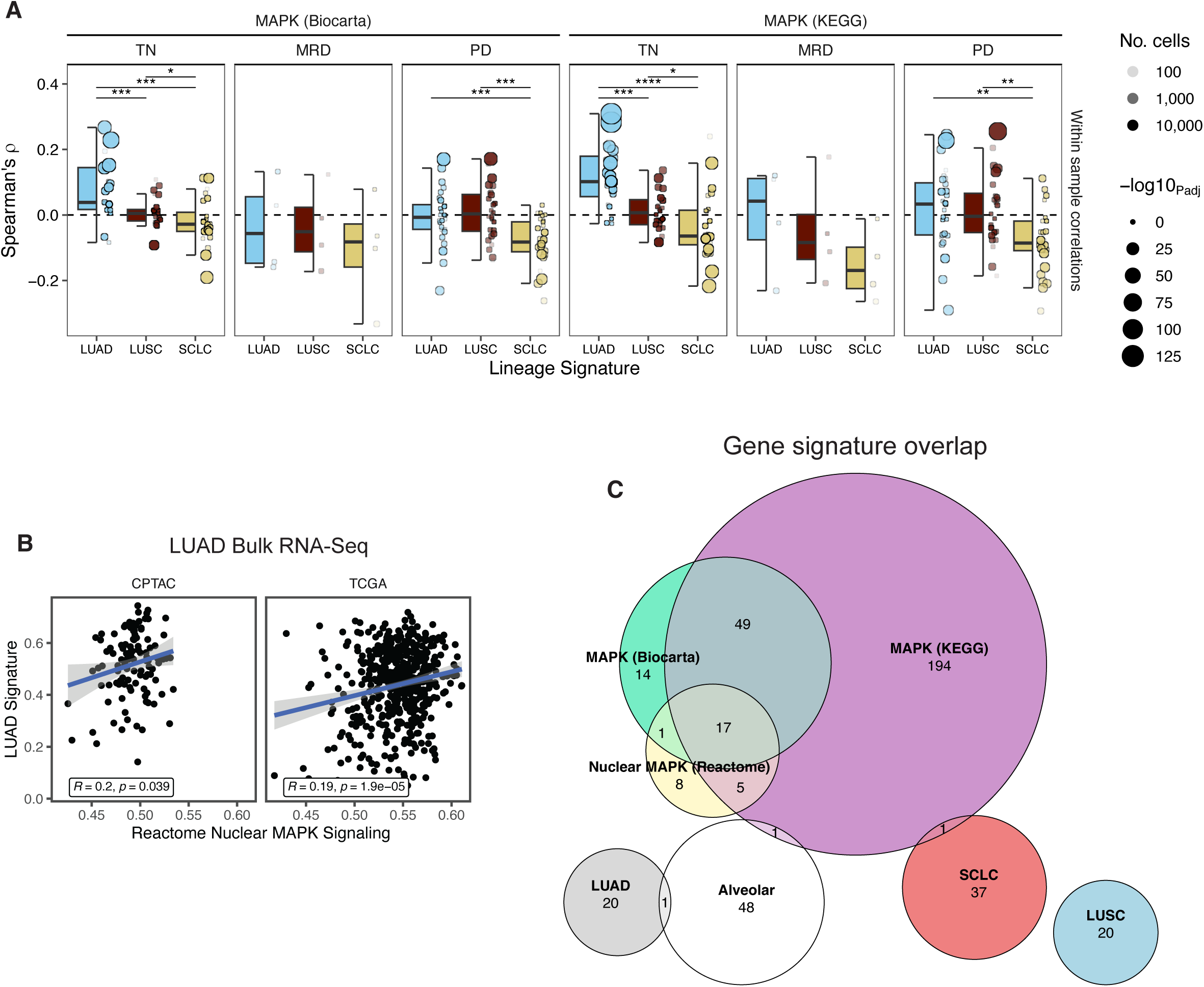
Lineage signatures and MAPK pathway activity. **A.** Box and dotplots showing within sample correlations between the Biocarta and KEGG MAPK pathways and LUAD, LUSC and SCLC signatures respectively. Each dot corresponds to a correlation test performed across tumor cells within an individual sample, sized by the level of significance (holm-adjusted), and shaded based on total cell counts. The y-axis indicates Spearman’s ρ. Significance levels of Wilcoxon rank-sum tests comparing the distributions of correlations are encoded as *’: p<0.05, ‘**’: p<0.01 ‘***’: p<0.001, ‘****’: p<0.0001. **B.** Scatterplots of LUAD and Reactome Nuclear MAPK signatures in the TCGA and CPTAC LUAD bulk RNA-seq datasets. **C.** Euler plot showing the overlap in genesets.

**Supplementary Figure 9.**
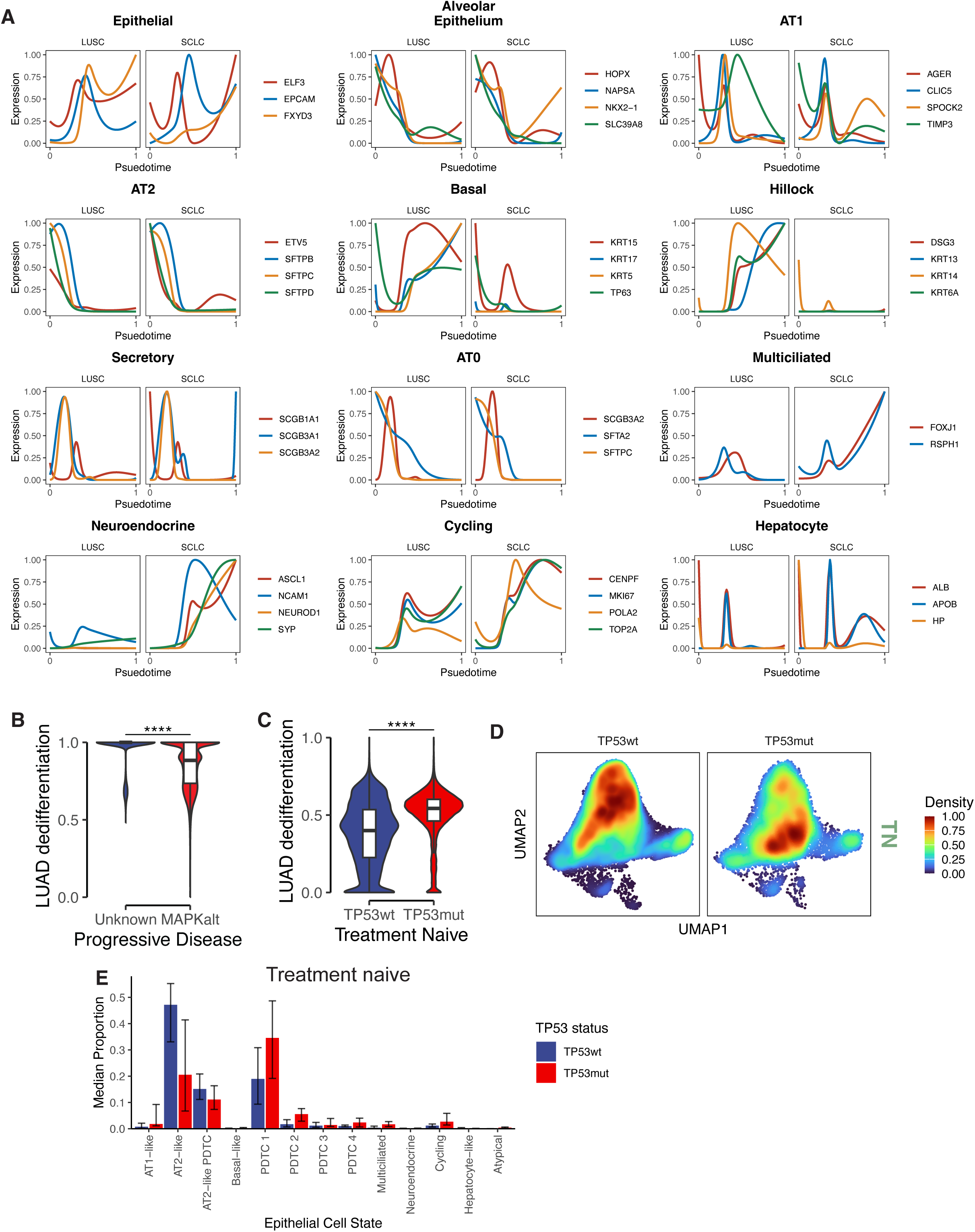
Histotime analysis. **A.** Scaled expression of epithelial lineage markers (highlighted in Fig. 3B) in the LUSC and SCLC histotime branches respectively. **B.** Histotime derived LUAD dedifferentiation scores in MAPK altered versus not progressive disease tumor cells. Wilcoxon rank-sum test significance is encoded as ‘****’: p<0.0001. **C.** Histotime derived LUAD dedifferentiation scores in *TP53* mutant versus wildtype treatment-naive tumor cells. Wilcoxon rank-sum test significance is encoded as ‘****’: p<0.0001. **D.** Density of tumor cells from *TP53* mutant and wildtype samples overlaid on the epithelial cell UMAP from Fig. 3A. **E.** Median proportions of *TP53* mutant and wildtype tumor cells belonging to each epithelial cell state. Error bars indicate interquartile range.

## TABLES

**Table S1** Cohort metadata

**Table S2** MSK-IMPACT somatic mutation table

**Table S3** SCLC signature genes

**Table S4** Features used as input for histotime analysis

